# Integrated Single-Cell and Spatial Transcriptomic Analysis Reveals an Aging-Associated Fibroblast Subtype Linked to Tumor Progression in Human Skin

**DOI:** 10.1101/2025.07.09.663827

**Authors:** Haibin Wu, Danping Pan

## Abstract

Aging is a major risk factor for the development of many cancers, yet the mechanisms underlying this increased susceptibility remain incompletely understood. Traditionally, the accumulation of genetic mutations over time has been considered a primary driver of age-related tumorigenesis. However, emerging evidence highlights the critical role of the aging tissue microenvironment—including changes in immune, stromal, and epithelial compartments—in shaping cancer initiation and progression. In this study, we employed integrated single-cell and spatial transcriptomics to systematically characterize age-associated alterations in human skin and skin cancers. Our analysis uncovered widespread, cell type-specific transcriptional reprogramming with age, revealing key pathways and cellular populations that may contribute to a pro-tumorigenic environment. Notably, we identified a previously unrecognized fibroblast subtype marked by SFRP2 expression, which expands with age and is associated with poor prognosis in basal cell carcinoma. These fibroblasts appear to enhance Wnt signaling within the tumor niche, suggesting a potential mechanism by which the aging stroma supports malignancy. Collectively, our findings shed light on how age-related changes in tissue ecosystems may predispose to cancer and point to novel therapeutic opportunities for targeting the aged tumor microenvironment.

## INTRODUCTION

Skin is the largest organ of the body, serving as a critical barrier that protects against dehydration, ultraviolet (UV) radiation, and invasion by pathogens(1, 2). It is a complex organ composed of diverse cell types, including keratinocytes, fibroblasts, melanocytes, and immune cells, all of which work in concert to maintain tissue homeostasis. However, aging profoundly alters the structural, cellular, and molecular composition of the skin, diminishing its regenerative capacity and resilience. Aging is also a well-established risk factor for a variety of diseases, including cancer(3, 4).

Skin cancer is among the most prevalent malignancies in the aging population, with basal cell carcinoma (BCC), squamous cell carcinoma (SCC), and melanoma being the most common types. Although accumulated genetic mutations have historically been considered the primary drivers of cancer development, increasing evidence highlights the pivotal role of the tumor microenvironment (TME) in cancer initiation and progression [4]. The TME comprises stromal and immune cells that interact dynamically with cancer cells, influencing tumor growth, invasion, and response to therapy. In aging individuals, the TME undergoes significant alterations, including chronic inflammation, extracellular matrix (ECM) remodeling, and immunosenescence, all of which contribute to the increased susceptibility to skin cancer(5).

Immunosenescence, the gradual deterioration of the immune system with age, is a critical factor in the failure of the immune system to recognize and eliminate tumor cells effectively. This age-associated decline includes reduced antigen presentation, diminished T-cell activation, and the accumulation of immunosuppressive cells, such as regulatory T cells and myeloid-derived suppressor cells(4, 6, 7). These changes create an immune environment that is less effective at tumor surveillance and more permissive to cancer progression. Additionally, stromal cells, particularly fibroblasts, undergo phenotypic and functional changes during aging that reshape the ECM and influence the behavior of surrounding cells(5, 8). Emerging technologies, such as single-cell RNA sequencing (scRNA-seq) and spatial transcriptomics, now allow for unprecedented resolution in profiling cellular heterogeneity and gene expression changes in complex tissues(9, 10). These tools have illuminated age-related shifts in cellular composition and molecular pathways across a variety of tissues, including the skin(11). Despite these advances, our understanding of how aging impacts specific cell types within the skin and their roles in cancer progression remains incomplete.

In this study, we leverage integrated scRNA-seq and spatial transcriptomics datasets to construct a comprehensive atlas of aging-related changes in human skin. We focus on fibroblasts, a key stromal cell type that plays a central role in maintaining skin integrity and modulating the TME. Our analyses reveal a novel fibroblast subtype, characterized by high expression of Secreted Frizzled-Related Protein 2 (SFRP2), which becomes more abundant with aging and is strongly associated with poor prognosis in BCC. SFRP2-positive fibroblasts activate Wnt signaling pathways, promoting tumor progression and creating a pro-tumorigenic microenvironment. By elucidating the cellular and molecular mechanisms underlying age-related changes in the skin, this work provides novel insights into the interplay between aging, fibroblast function, and cancer vulnerability. Our findings underscore the importance of targeting aging-associated pathways as a therapeutic strategy for age-related cancers and highlight SFRP2-positive fibroblasts as a potential biomarker and therapeutic target.

## METHOD DETAILS

### Data Acquisition and Preprocessing

Publicly available single-cell RNA sequencing (scRNA-seq) datasets were downloaded from the NCBI Sequence Read Archive (SRA) and Genome Sequence Archive (GSA) using sratoolkit (v2.11.2) or Aspera Connect (v4.1.1) for high-speed transfer(12). The dataset comprised skin samples from 14 female donors aged 18 to 76 years, previously published under GEO accession number GSE130973(13) and GSA accession number HRA000395(14). These datasets were merged, integrated, and processed for downstream analysis. Human melanoma scRNA-seq data were obtained from GEO accession GSE115978(15). For squamous cell carcinoma (SCC), both scRNA-seq and spatial transcriptomics data were downloaded from GEO under accessions GSE218170(16) and GSE208253(17), respectively. For basal cell carcinoma (BCC), scRNA-seq and spatial transcriptomics data were retrieved from ArrayExpress under accessions E-MTAB-13085 and E-MTAB-13084(18).For scRNA-seq datasets, the FASTQ files were aligned to the GRCh38 human genome reference using the CellRanger pipeline (v6.1.2). The resulting gene expression matrices were aggregated into a single matrix for downstream analysis. Cells with fewer than 200 detected genes, mitochondrial gene content exceeding 10%, or total gene counts greater than 2.5 standard deviations above the mean were excluded.

### Visium data processing

Raw Visium spatial transcriptomics data (BCL and FASTQ files) were processed using the SpaceRanger v1.3.1 pipeline (10x Genomics). For each sample, gene-barcode matrices were loaded into R and organized using the SummarizedExperiment and SingleCellExperiment frameworks to retain spatial metadata, including histological image alignment and tissue position coordinates. Spots were initially filtered based on quality control metrics using the perCellQCMetrics function from the scran v1.14.3 package(19). Specifically, spots were excluded if they contained fewer than 200 detected genes, fewer than 500 total UMI counts, or >15% mitochondrial gene content. After quality filtering, gene counts were normalized using scran’s computeSumFactors, followed by log-transformation with logNormCounts from scater v1.14.3(20). Cell type annotation

Cell types were annotated using a combination of automated clustering and manual curation. Initial clustering was performed with Seurat’s FindClusters function using a resolution of 0.5. Cluster identities were assigned based on canonical marker gene expression, including KRT14 (keratinocytes), COL1A1 (fibroblasts), PECAM1 (endothelial cells), and CD3D (T cells). Differentially expressed genes (DEGs) were identified using the Wilcoxon rank-sum test, with p-values adjusted for multiple testing using the Benjamini-Hochberg method. Marker genes were further validated by cross-referencing published datasets(13, 14).

### Trajectories analysis

Single-cell trajectory analysis was performed using Monocle3(21) to infer developmental trajectories of fibroblasts, enabling the identification of transitional states leading to SFRP2-positive fibroblasts. The pseudotime dynamics of aging-associated gene expression were visualized to assess how fibroblast subtypes evolve with age.

### Ligand-Receptor interaction analysis

Cellular communication networks were analyzed using the CellChat package(22), focusing on ligand-receptor interactions mediated by fibroblasts. Pathway enrichment analysis identified key signaling axes. The signaling strength was quantified and visualized as circular plots for young, middle-aged, and old skin samples.

### Prognostic analysis

Prognostic implications of aging-associated gene programs were assessed using The Cancer Genome Atlas (TCGA) datasets for BCC, melanoma, and squamous cell carcinoma (SCC). Patients were stratified into high and low expression groups based on aging-related gene signatures expression, and Kaplan-Meier survival analysis was performed. Hazard ratios and statistical significance were calculated using the survival package in R.

### Statistical Analysis

Statistical analyses were conducted in R (v4.2.2) and Python (v3.9). DEGs were identified with a threshold of adjusted p-value <0.05 and absolute log2 fold-change >0.25. Batch correction and integration metrics were calculated using the scIB framework. All plots, including UMAPs, heatmaps, and spatial transcriptomics maps, were generated using ggplot2, ComplexHeatmap, and Seurat visualization tools.

## RESULTS

### Construction of an Integrated Immune Cell Atlas of Human Skin Aging

To elucidate the landscape of immune cell dynamics during skin aging, we integrated two publicly available single-cell RNA sequencing (scRNA-seq) datasets spanning 14 female donors aged 18 to 76 years. After stringent quality control, a total of 216,778 high-quality cells were retained for analysis. We benchmarked several data integration methods (Harmony, LIGER, Scanorama, and Seurat) using 14 performance metrics assessing batch effect correction and biological variance retention. Harmony outperformed other methods and was selected for further analysis (Figure 1A).

**Figure 1.**
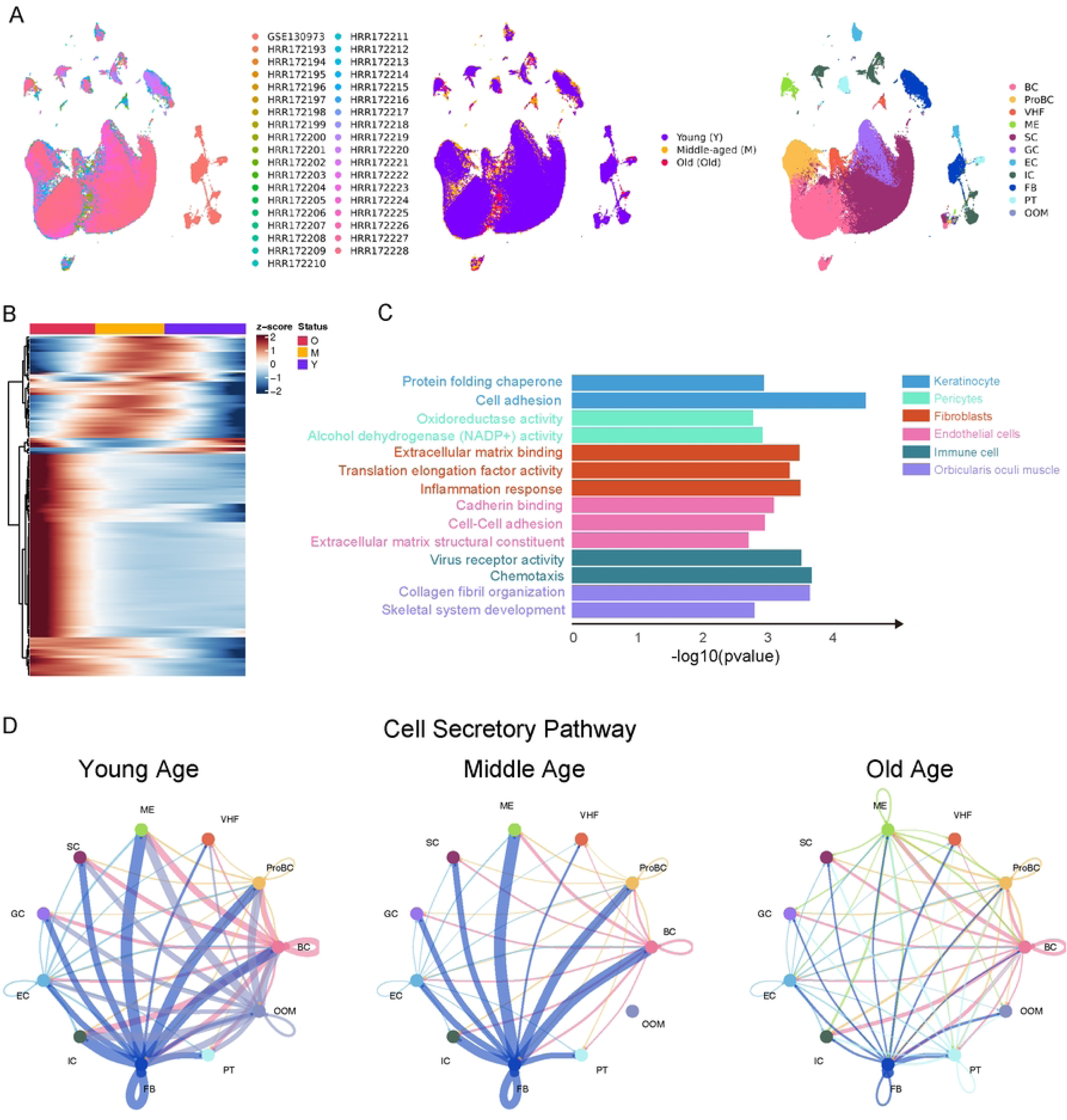
Single-cell RNA sequencing atlas of human aging. (A) UMAP plot showing integrated single-cell RNA-seq data from human skin across ages. Clusters represent major skin cell types, including BC, VHF, ME, SC, GC, EC, IC, FB, and LC, highlighting age-associated shifts in cellular composition. (B) Heatmap of age-related gene expression across skin cell types. Color scale (blue to red) reflects expression levels of genes significantly altered with aging, emphasizing cell type-specific transcriptional changes. (C) Enriched GO terms in aging skin cells. Bar plots show top GO terms for six cell types with the most age-related transcriptional changes, revealing key biological processes such as ECM organization and immune response. (D) CellChat analysis of fibroblast signaling across age groups (young, middle, old). Circular plots depict evolving fibroblast communication dynamics, highlighting age-related shifts in key signaling pathways within the skin microenvironment.

Following integration, Uniform Manifold Approximation and Projection (UMAP) visualization revealed clear clustering of major immune cell populations, including T cells, NK cells, B cells, macrophages, dendritic cells (DCs), and mast cells. These clusters were consistent across age groups, confirming the robustness of integration and annotation. Notably, the distribution and density of specific immune subsets shifted with aging, suggesting age-associated alterations in the immune landscape.

To further explore molecular aging signatures, we performed differential gene expression analysis across immune cell types, followed by hierarchical clustering. The resulting heatmap revealed distinct aging-associated transcriptional programs, particularly in T cells and macrophages (Figure 1B). Gene ontology (GO) enrichment analysis of differentially expressed genes highlighted pathways involved in immune cell activation, extracellular matrix organization, and apoptotic signaling (Figure 1C), implicating both immunosenescence and stromal remodeling processes.

We next assessed intercellular communication dynamics using CellChat. Network analyses of signaling interactions among immune cells in young, middle-aged, and old skin demonstrated progressive rewiring of communication networks with age (Figure 1D–F). Aging was associated with increased signaling complexity, particularly involving fibroblast–immune and macrophage–T cell interactions, with elevated activity in pathways such as TNF, CXCL, and MIF. These findings suggest that immune aging in skin is not only marked by intrinsic transcriptomic changes but also by altered cellular crosstalk within the tissue microenvironment.

### Cell-type-specific aging programs are activated in distinct skin cancers and associate with patient prognosis

To investigate how aging impacts skin cancer at the cellular level, we analyzed single-cell RNA sequencing data from melanoma, squamous cell carcinoma (SCC), and basal cell carcinoma (BCC) derived from public datasets. UMAP visualizations revealed a diverse cellular landscape in each cancer type, including immune cells, endothelial cells, fibroblasts, and keratinocytes (Figure 2A–C). We applied a curated aging gene signature to evaluate the activation of aging programs across these cell populations.

**Figure 2.**
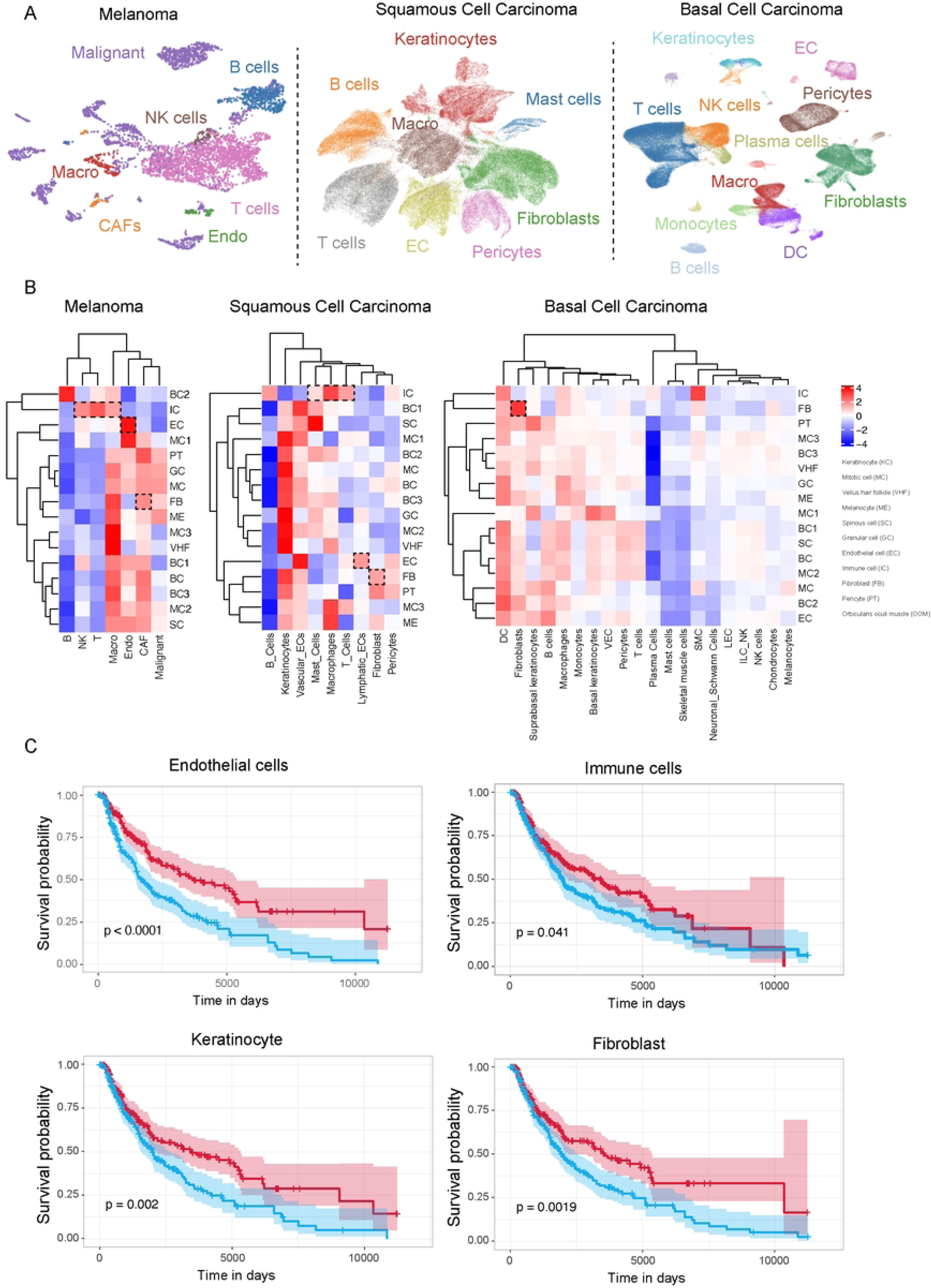
Aging-related niches associated with poor prognosis in skin cancer. (A) UMAP plots of single-cell profiles from melanoma, SCC, and BCC samples, showing diverse tumor microenvironment cell types, including CAFs, immune cells, endothelial cells, and fibroblasts. (B) Heatmaps of aging signature scores across cell types in skin cancers. Color scale (blue to red) indicates expression levels of aging-related genes, revealing cell type-specific aging effects within tumors. (C) Kaplan–Meier survival curves linking aging signatures in specific cell types to patient outcomes. Higher aging signature expression correlates with poorer survival, highlighting the prognostic impact of age-related changes in the tumor microenvironment.

Our analysis uncovered strong cell-type-specific aging patterns within and across tumor types. In melanoma, aging programs were prominently upregulated in fibroblasts, endothelial cells, and immune cells (Figure 2D). In contrast, BCC showed more restricted aging signatures, with minimal activation in immune cells but notable aging-associated transcriptional changes in fibroblasts and endothelial cells. SCC displayed an intermediate profile, with clear aging program activation in both fibroblasts and immune compartments (Figure 2E–F). These findings highlight both shared and distinct aging trajectories among cell types and tumor contexts.

To assess the clinical relevance of these aging programs, we examined their association with patient outcomes using bulk RNA-seq and clinical data from The Cancer Genome Atlas (TCGA). Kaplan–Meier survival analyses revealed that higher expression of aging-related signatures in endothelial cells, fibroblasts, keratinocytes, and immune cells was significantly associated with poorer prognosis in melanoma patients (Figure 2G–J). These results suggest that aging-related molecular changes within specific stromal and immune compartments may actively shape tumor behavior and impact clinical outcomes.

Furthermore, the conserved activation of fibroblast and endothelial cell aging across all three skin cancer types suggests a shared mechanism by which the aged microenvironment supports tumor progression. This is consistent with previous studies demonstrating that fibroblast-mediated extracellular matrix (ECM) remodeling—driven by matrix metalloproteinases (MMPs) such as MMP-2, MMP-9, and MMP-14—facilitates tumor invasion and metastasis. Similarly, aged endothelial cells contribute to tumor vascularization and immune modulation via increased expression of pro-angiogenic factors such as VEGF and FGF.

In contrast, immune cell aging was much less evident in BCC, underscoring the importance of tumor-type-specific aging profiles. These findings emphasize the need for precision therapeutic strategies that consider not only tumor mutations but also the aging status of distinct stromal and immune components within the tumor microenvironment.

### SFRP2⁺ Fibroblasts Spatially Associate with Tumor Regions and Activate Wnt Signaling in BCC

To explore the role of aging-associated fibroblasts in basal cell carcinoma (BCC), we analyzed the spatial organization and signaling profiles of fibroblasts using integrated single-cell and spatial transcriptomics. Spatial proximity analysis revealed strong clustering of tumor and stromal compartments, particularly in localized hotspots within the tissue (Figure 3A). These spatial interactions suggest a compartmentalized microenvironment where tumor and fibroblast zones closely associate.

**Figure 3.**
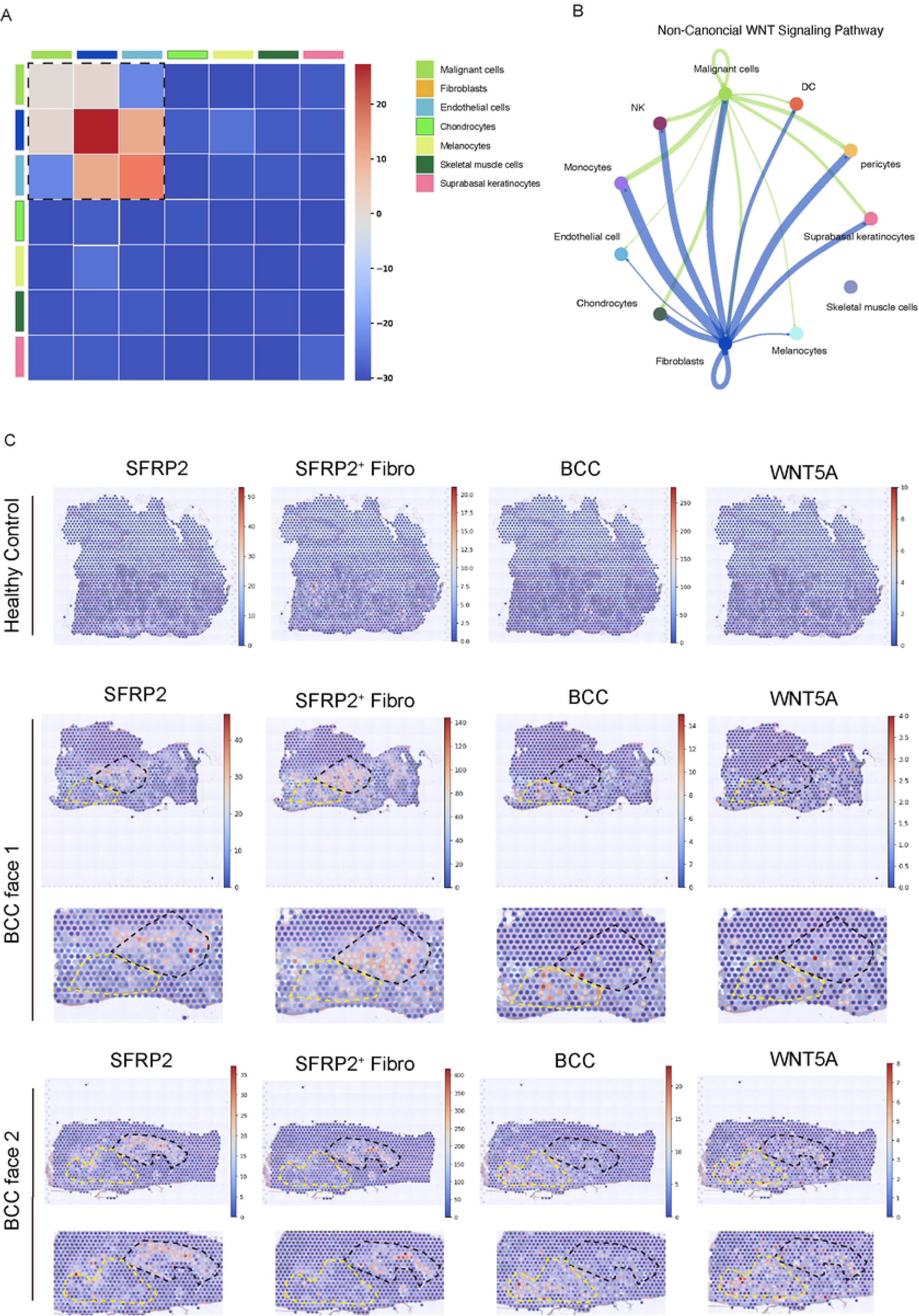
SFRP2+ fibroblasts promote Wnt signaling in basal cell carcinomas (BCCs). (A) Heatmap showing spatial proximity among BCC spatial transcriptomic spots, with red indicating closely clustered tumor and stromal regions. (B) Ligand-receptor interaction network from BCC scRNA-seq data. Line thickness reflects interaction strength, highlighting a prominent SFRP2–LRP5/6 axis driven by fibroblasts and linked to Wnt signaling activation. (C) Spatial maps of SFRP2⁺ fibroblasts, BCC cells, and WNT5A expression. Co-localization of fibroblasts and Wnt pathway activity in tumor areas suggests a fibroblast-mediated pro-tumorigenic niche.

We next investigated cell–cell signaling interactions using ligand–receptor inference on BCC single-cell RNA-seq data. As shown in Figure 3B, fibroblasts emerged as central signaling hubs, with a strong interaction between fibroblast-derived SFRP2 and its receptors LRP5/6 expressed in neighboring stromal and tumor cells. The prominence of this axis suggests that SFRP2 may function as a key modulator of Wnt signaling activation in the tumor microenvironment.

To spatially validate these interactions, we mapped the expression of SFRP2, BCC tumor markers, and WNT5A across multiple tissue sections (Figure 3C). SFRP2⁺ fibroblasts were enriched in the tumor-surrounding stroma, while WNT5A expression localized to overlapping regions within and around the tumor core. The repeated co-localization of SFRP2 and WNT ligands across tissue depths supports the conclusion that SFRP2⁺ fibroblasts contribute to a pro-tumorigenic niche by enhancing non-canonical Wnt signaling within BCC lesions.

### Aged Fibroblasts Activate Wnt Signaling in SCC Tumor-Stroma Interfaces

To assess whether similar fibroblast-mediated mechanisms operate in squamous cell carcinoma (SCC), we analyzed Visium spatial transcriptomics from SCC samples. UMAP projection of spatial spots revealed distinct grouping by donor origin (Figure 4A), tumor compartment (core, edge, surrounding; Figure 4B), and aggregated tumor zones (Figure 4C), confirming spatial heterogeneity across samples.

**Figure 4.**
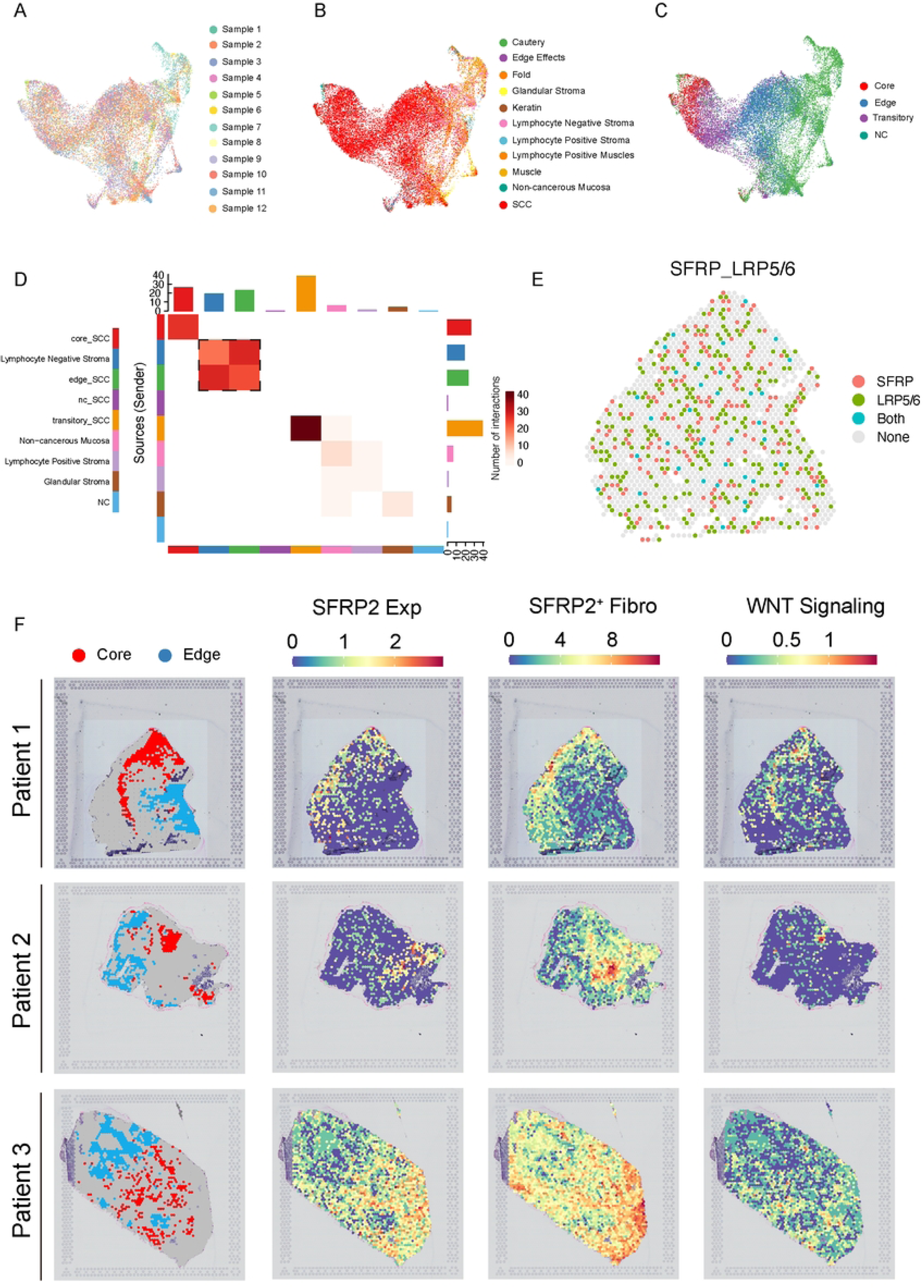
SFRP2+ fibroblasts promote Wnt signaling in the tumor microenvironment of SCC. (A–C) UMAP plots of Visium spatial spots from SCC samples. (A) Spots colored by sample origin. (B) Spots labeled by spatial zones (core, edge, surrounding). (C) Grouped zones highlight transcriptomic patterns at tumor boundaries. (D) Heatmap showing spatial proximity among tumor zones. Higher intensity indicates closer spatial interaction, illustrating SCC microenvironment organization. (E) Ligand–receptor analysis identifies fibroblast-derived SFRP2 interacting with LRP5/6, suggesting Wnt signaling activation. (F) Spatial maps of SFRP2⁺ fibroblasts and Wnt signaling scores in SCC. Enrichment in tumor cores and interfaces points to fibroblast-driven modulation of the tumor niche.

**Figure 5.**
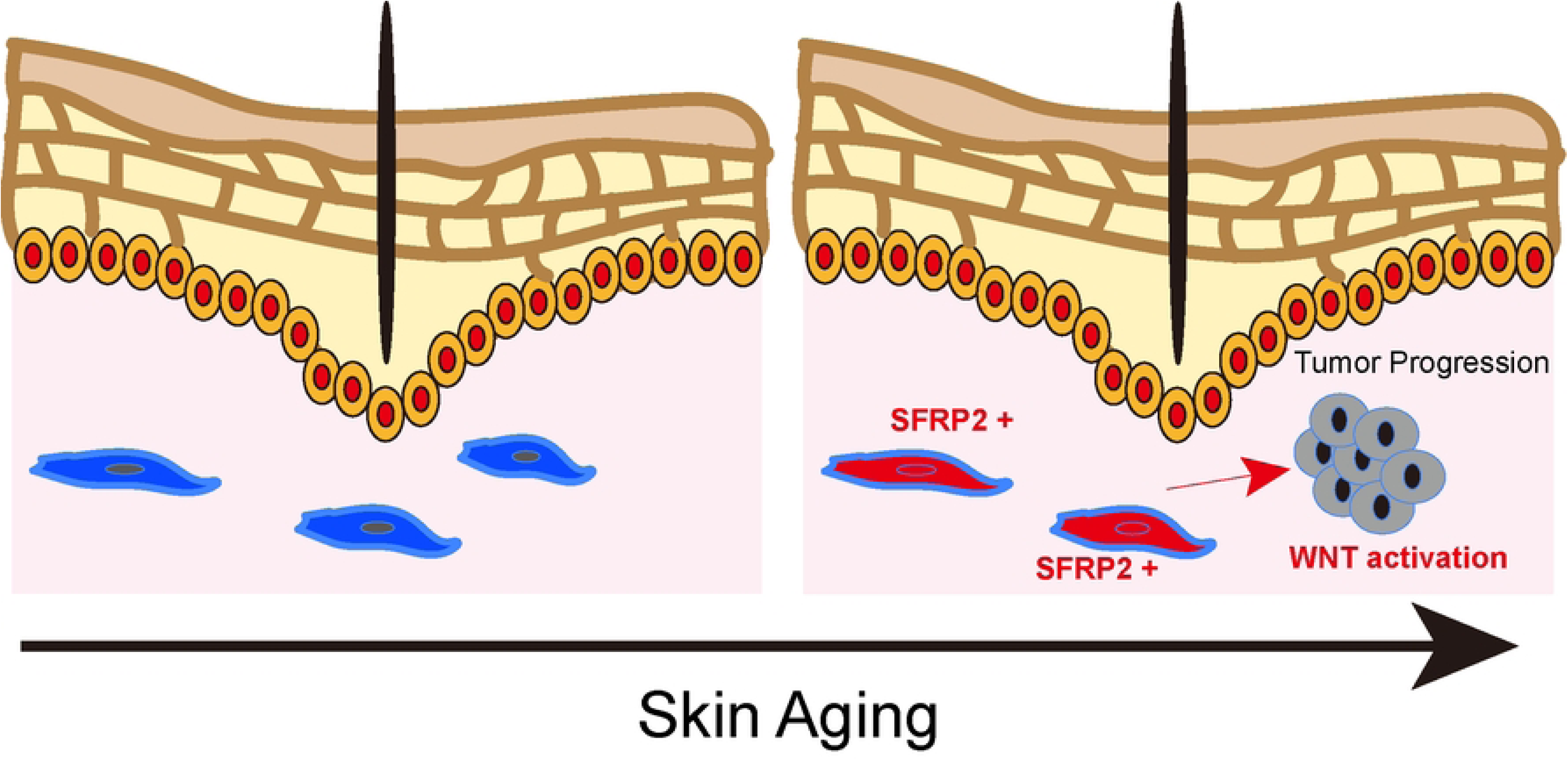
Diagram illustrating how aging promotes fibroblast transitions and increases tumor vulnerability. Schematic illustrating aging-induced transition of fibroblasts into SFRP2⁺ fibroblasts. Left: In young skin, fibroblasts maintain tissue homeostasis. Right: With aging, fibroblasts shift to an SFRP2⁺ state, activating Wnt signaling and fostering a pro-tumorigenic environment, increasing cancer susceptibility.

A spatial proximity matrix showed strong connectivity between tumor cores and adjacent stromal regions (Figure 4D), indicating active tumor–stroma interactions. Ligand–receptor analysis again identified SFRP2–LRP5/6 interactions, with fibroblasts expressing the ligand and tumor-adjacent cells expressing corresponding receptors (Figure 4E). These findings indicate that Wnt pathway activation through aging-associated fibroblast signaling is a conserved feature in SCC, although the spatial pattern appeared broader and less centralized compared to BCC.

## DISCUSSION

Aging is increasingly recognized as a key contributor to cancer susceptibility, not only through the accumulation of genetic mutations but also via profound changes in the tissue microenvironment(23, 24). In this study, we integrated single-cell and spatial transcriptomics to dissect age-associated alterations in human skin and skin cancers. Our results highlight marked shifts in cellular composition and gene expression programs with age, particularly implicating fibroblasts as central players in creating a tumor-permissive environment.

Among these, a previously uncharacterized population of SFRP2-positive fibroblasts emerged as significantly enriched in aged skin and further expanded in basal cell carcinoma (BCC). Although traditionally viewed as Wnt antagonists, SFRP2 has been shown to activate non-canonical Wnt/Ca²⁺ signaling in cancer contexts and to promote tumor angiogenesis via Fzd5/NFAT pathways(25, 26). Our data support and extend this view, suggesting that SFRP2⁺ fibroblasts contribute to tumor progression by promoting Wnt signaling, extracellular matrix remodeling, and angiogenesis—hallmarks of a pro-tumorigenic niche. This aligns with prior findings that fibroblast-derived factors, including MMPs and VEGF, drive tumor invasion and metastasis. Importantly, we found that aging signatures varied considerably across cell types. While fibroblast aging was prominent in BCC, immune and endothelial compartments displayed distinct aging trajectories. For example, endothelial cells in aged skin exhibited elevated angiogenic programs, including VEGF signaling, known to enhance tumor vascularization. In contrast, immune aging was more pronounced in melanoma and squamous cell carcinoma (SCC), consistent with the higher immune cell infiltration and immunosuppressive microenvironments observed in those cancers(27). These differences reinforce the need for tumor type-specific strategies when targeting aging-associated pathways.

Spatial transcriptomics was essential for resolving cell localization and interaction networks, enabling us to map the enrichment and spatial clustering of SFRP2-positive fibroblasts near tumor sites. This integrative approach helped overcome limitations of single-cell data in estimating abundance and provided a contextual understanding of how aging fibroblasts interact with surrounding cells to influence cancer behavior. Therapeutically, targeting SFRP2-positive fibroblasts or modulating aberrant Wnt signaling may offer novel strategies for age-associated skin cancers(27, 28). SFRP2 itself may serve as a biomarker to stratify patients who might benefit from such interventions. More broadly, this study underscores the importance of incorporating aging biology into cancer research frameworks. Future studies should investigate whether similar fibroblast-driven mechanisms operate in other aging tissues and cancer types, and explore interventions that mitigate age-induced stromal changes to improve cancer outcomes.

## ACKNOWLEDGMENTS

We thank BioYishang for providing writing guidance and support during the preparation of this manuscript.

